# Uncharacterized bacterial structures revealed by electron cryotomography

**DOI:** 10.1101/108191

**Authors:** Megan J. Dobro, Catherine M. Oikonomou, Aidan Piper, John Cohen, Kylie Guo, Taylor Jensen, Jahan Tadayon, Joseph Donermeyer, Yeram Park, Benjamin A. Solis, Andreas Kjær, Andrew I. Jewett, Alasdair W. McDowall, Songye Chen, Yi-Wei Chang, Jian Shi, Poorna Subramanian, Cristina V. Iancu, Zhuo Li, Ariane Briegel, Elitza I. Tocheva, Martin Pilhofer, Grant J. Jensen

## Abstract

**SUMMARY STATEMENT:** Here we present a survey of previously uncharacterized structures we have observed in bacterial cells by electron cryotomography, in the hopes of spurring their identification and study.

**ABSTRACT:** Electron cryotomography (ECT) can reveal the native structure and arrangement of macromolecular complexes inside intact cells. This technique has greatly advanced our understanding of the ultrastructure of bacterial cells. Rather than undifferentiated bags of enzymes, we now view bacteria as structurally complex assemblies of macromolecular machines. To date, our group has applied ECT to nearly 90 different bacterial species, collecting more than 15,000 cryotomograms. In addition to known structures, we have observed several, to our knowledge, uncharacterized features in these tomograms. Some are completely novel structures; others expand the features or species range of known structure types. Here we present a survey of these uncharacterized bacterial structures in the hopes of accelerating their identification and study, and furthering our understanding of the structural complexity of bacterial cells.

## INTRODUCTION

The history of cell biology has been punctuated by advances in imaging technology. In particular, the development of electron microscopy in the 1930s produced a wealth of new information about the ultrastructure of cells (Ruska, 1987). For the first time, the structure of cell envelopes, internal organelles, cytoskeletal filaments and even large macromolecular complexes like ribosomes became visible. A further advance came in the 1980s and 1990s with the development of electron cryotomography (ECT) (Koster et al., 1997), which allows small cells to be imaged intact in 3D in a near-native, “frozen-hydrated” state to “macromolecular” (~4 nm) resolution, without the limitations and artifacts of more traditional specimen preparation methods (Pilhofer et al., 2010).

ECT has helped reveal the previously unappreciated complexity of “simple” bacterial cells. Our group has been using ECT to study bacteria for more than a decade, generating more than 15,000 tomograms of 88 different species. These tomograms have revealed new insights into, among other things, the bacterial cytoskeleton (Komeili et al., 2006, Li et al., 2007, Pilhofer et al., 2011, Swulius & Jensen, 2012), cell wall architecture (Gan et al., 2008, Beeby et al., 2013), morphogenesis (Ebersbach et al., 2008), metabolism (Iancu et al., 2007), motility (Murphy et al., 2006, Chen et al., 2011, Abrusci et al., 2013, Chang et al., 2016), chemotaxis (Briegel et al., 2012), sporulation (Tocheva et al., 2011), cell-cell interactions (Basler et al., 2012), and phage infection (Guerrero-Ferreira et al., 2011) (for a summary with more references from our and others’ work, see (Oikonomou & Jensen, 2016)).

A major hurdle in such studies is identifying the novel structures observed in tomograms. In some cases, we have identified structures by perturbing the abundance (either by knockout or overexpression) of candidate proteins (Ingerson-Mahar et al., 2010). In others, we have used correlated light and electron microscopy (CLEM) to locate tagged proteins of interest (Briegel et al., 2008, Chang et al., 2014). In one striking example, we observed 12- and 15-nm tubes in our tomograms of *Vibrio cholerae* cells. Ultimately, in collaboration with John Mekalanos’ group, we identified them as type VI secretion systems (T6SS), which immediately led to the insight that the bacterial T6SS functions as a phage-tail-like, contractile molecular dagger (Basler et al., 2012).

Many other novel structures we have observed, though, remain unidentified. In some cases, we have published papers describing the novel structures seen in a particular species (e.g. (Murphy et al., 2008, Muller et al., 2014)), but many have never been published. We therefore conducted a visual survey of the tomograms collected by our group, curated in the Caltech Tomography Database (Ding et al., 2015), as of 2015 and present here a catalog of previously undescribed bacterial structures. Some structures are, to our knowledge, completely novel; others belong to known types but present additional features or an expanded species range. We hope that sharing these images will help spur their identification and study, contributing to our expanding understanding of bacterial cell biology. In addition, we look forward to a future in which custom microbes are designed for diverse medical and industrial purposes; an expanded “parts list” of structures to be repurposed will aid in this effort.

## RESULTS AND DISCUSSION

We performed a visual inspection of approximately 15,000 tomograms of intact, frozen-hydrated cells belonging to 88 species and identified what we believed to be novel structures. A summary of the results of this survey is shown in Supplementary Table 1, with features observed, species range, and frequency listed for each structure type. For full tomographic (3D) views of each feature, please see the accompanying supplementary movies at the following link: https://figshare.com/s/782461843c3150d27cfa.

### Extracellular structures

#### External appendages

In several tomograms of *Prosthecobacter debontii* we observed novel extracellular appendages, apparently attached to the cell membrane. Individual cells displayed up to 30 such appendages, which exhibited consistent size (~20 nm wide and ~50 nm long) and shape (Figure 1A). Subtomogram averaging of 105 particles revealed a distinctive structure: extending outward from the cell membrane, five legs were attached to a disc, which in turn connected to a smaller disc and a long neck region (Figure 1B,C). Individual particles showed that the structure culminated in two antenna-like filaments, which were likely averaged out in the average due to conformational variability. The appendages were observed in multiple cultures of the strain. While it remains unclear whether they originated intra- or extracellularly, no free-floating appendages were ever observed in the extracellular space. They may represent novel bacterial attachment organelles, a novel secretion system (though we would expect a cell envelope spanning complex) or a novel bacteriophage (though there is a notable lack of a capsid-like density).

**Figure 1.**
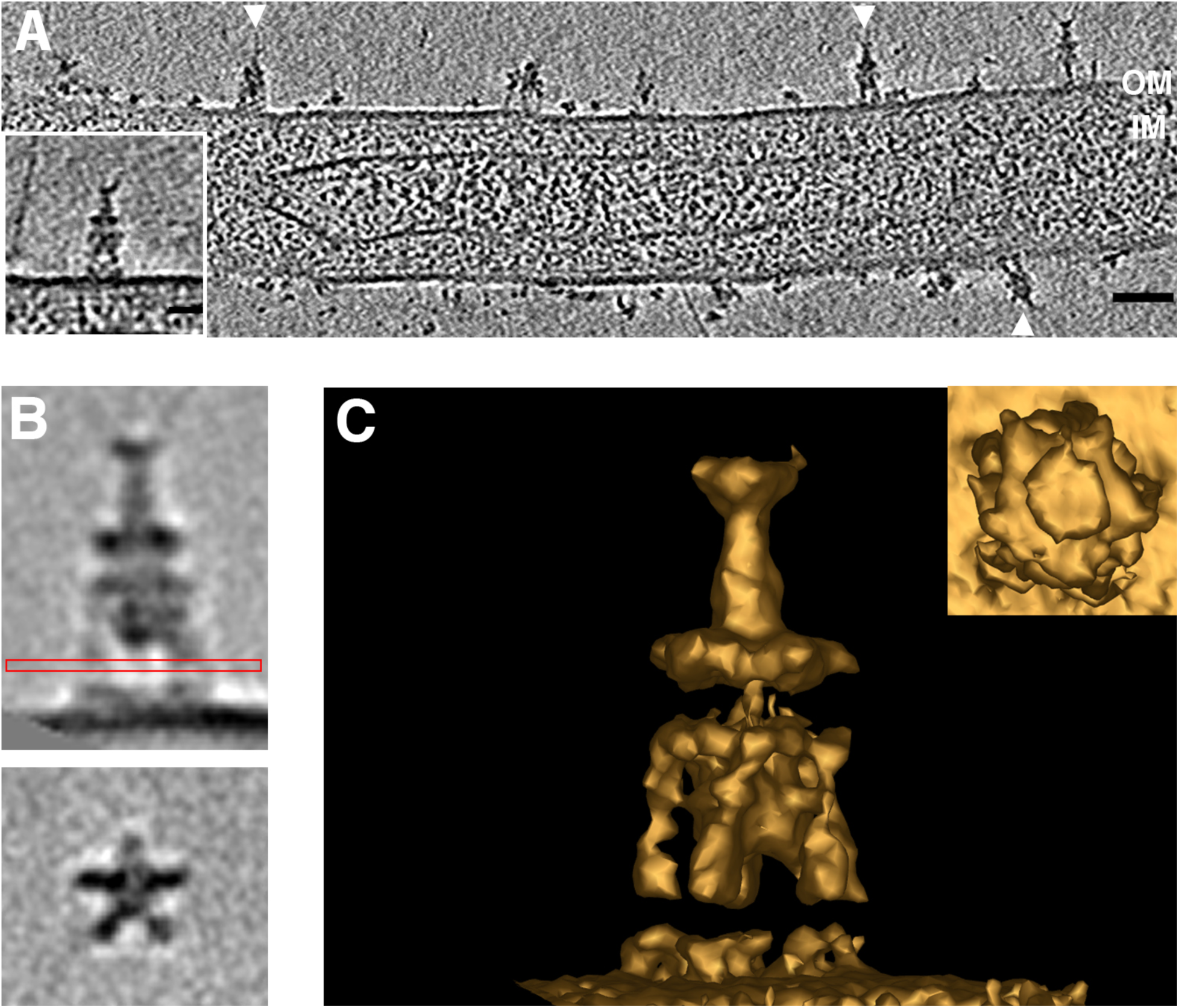
Novel *Prosthecobacter debontii* appendages. Multiple external appendages (arrowheads) were observed by ECT on *P. debontii* prosthecae (**A**). A central tomographic slice is shown, with a single appendage enlarged in the inset. Subtomogram averaging revealed the structure in more detail. Side (above) and top (below) views in (**B**) show the characteristic disc-like densities and the five legs attaching to the cell surface. The red box shows which view was used to rotate the image 90° for the bottom image. (**C**) shows a 3D isosurface of the average, seen from the side and top (inset). Scale bars 50 nm in (A) and 20 nm in inset.

We observed a different novel extracellular appendage in tomograms of *Azospirillum brasilense* cell poles. Thin hooks were seen extending out from the cell surface (Figure 2). Individual cells exhibited dozens of hooks, each ~3 nm wide and ~75 nm long, associated with the outer membrane. Hooks were seen in > 90% of wild-type cells as well as in a strain in which the operon encoding the Che1 chemotaxis system was deleted. They were seen in ~50 % of cells in which the Che4 chemotaxis system operon was deleted, and none were seen in cells lacking both the Che1 and Che4 operons. *A. brasilense* is a well-studied plant growth-promoting bacterium. Cells attach to plant roots through a two-step process (De Troch & Vanderleyden, 1996): a rapid, reversible adsorption thought to be mediated by the polar flagellum; and a slow, irreversible anchoring, thought to be mediated by an as-yet unidentified surface polysaccharide (Steenhoudt & Vanderleyden, 2000). A recent study reported that mutants in components of the Che4 chemotaxis system are defective in this root colonization (Mukherjee et al., 2016). *A. brasilense* cells also attach to conspecifics in the presence of elevated oxygen levels (Bible et al., 2015). Interestingly, it has been shown that mutants in components of the Che1 chemotaxis system form such attachments more rapidly than wild-type cells (Bible et al., 2012). The hooks we observed are vaguely reminiscent of the grappling hook-like structures that an archaeal species uses to anchor itself in biofilms (Moissl et al., 2005). It is therefore tempting to speculate that these hooks similarly play a role in adhesion, either to other *A. brasilense* cells or to plant roots.

**Figure 2.**
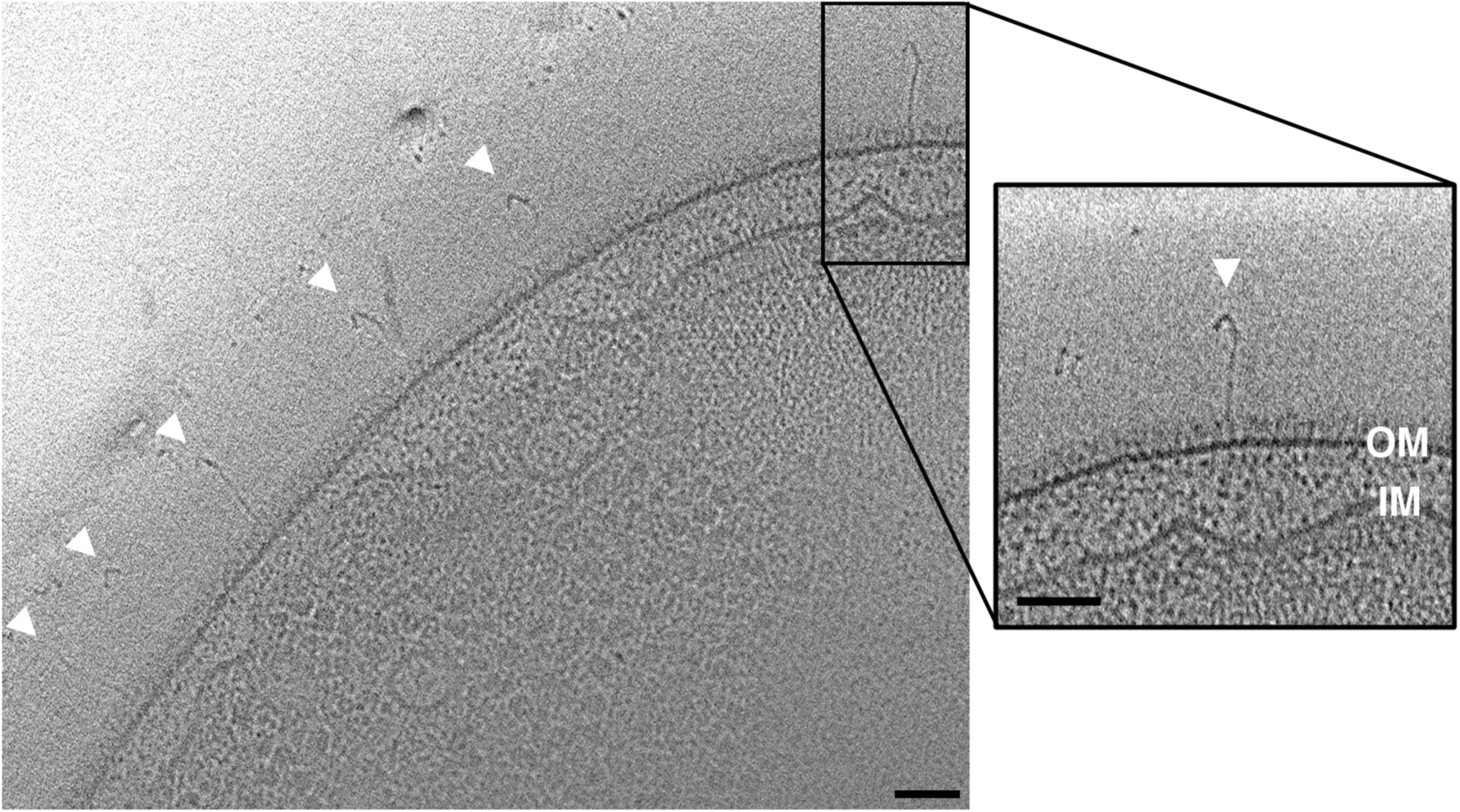
Novel *Azospirillum brasilense* hooks. Many hook-like structures were observed on the surface of *A. brasilense* cells. A central tomographic slice is shown, with arrowheads indicating hooks. A single hook is shown enlarged at right. Scale bar 50 nm.

In cells of strain JT5 (a bacterium isolated from termite gut and related to the *Dysgonomonas* genus), we observed abundant fimbriae concentrated at the cell poles (Figure 3). They were present in cells grown on cellulose or xylan, as well as in a condition inducing starvation. Their width (~4 nm), apparent flexibility, density on the cell envelope, and inhomogeneous distribution around the cell is consistent with curli, functional amyloids secreted by the type VIII secretion system that are involved in adhesion (Epstein et al., 2009, Van Gerven et al., 2015). Curli systems are relatively divergent at the sequence level, but are remarkably widespread phylogenetically, and the genes were reported to be present in Bacteroidetes (the phylum containing *Dysgonomonas*) (Dueholm et al., 2012). The appendages we observed in strain JT5 may therefore play a role in adhesion in the environment of the termite gut.

**Figure 3.**
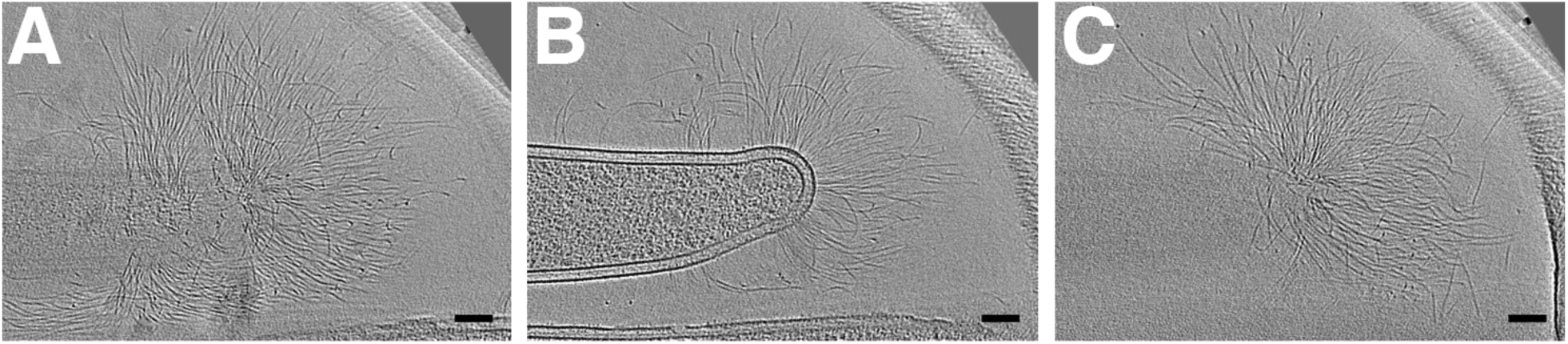
Strain JT5 fimbriae. (A-C) show slices at progressive z-heights through a cryotomogram of a cell of strain JT5 (related to the *Dysgonomonas* genus). Abundant fimbriae can be seen at the cell pole. Scale bars 100 nm.

#### Outer membrane vesicle chains

A wide variety of Gram-negative Bacteria produce outer membrane vesicles (Kulp & Kuehn, 2010). Recently, chained or tubular outer membrane structures have also been identified in a few species. In *Delftia* sp. Cs1-4, tubular sheaths called “nanopods” deliver outer membrane vesicles some distance from the cell (Shetty et al., 2011). In *Myxococcus xanthus*, chained extensions of the outer membrane transfer proteins and molecules between cells in a biofilm (Remis et al., 2014). In *Shewanella oneidensis*, nanowires used for extracellular electron transport can take the form of outer membrane vesicle chains (Pirbadian et al., 2014). We also observed chained extensions of the outer membrane in *Borrelia burgdorferi* strain B31 (Figure 4). Both spherical and tubular segments were observed in these chains. Outer membrane vesicles are important virulence factors in *B. burgdorferi* (Skare et al., 1995), and lipid exchange was recently observed between *B. burgdorferi* and host cells (Crowley et al., 2013); a chained vesicle morphology may aid in this process.

**Figure 4.**
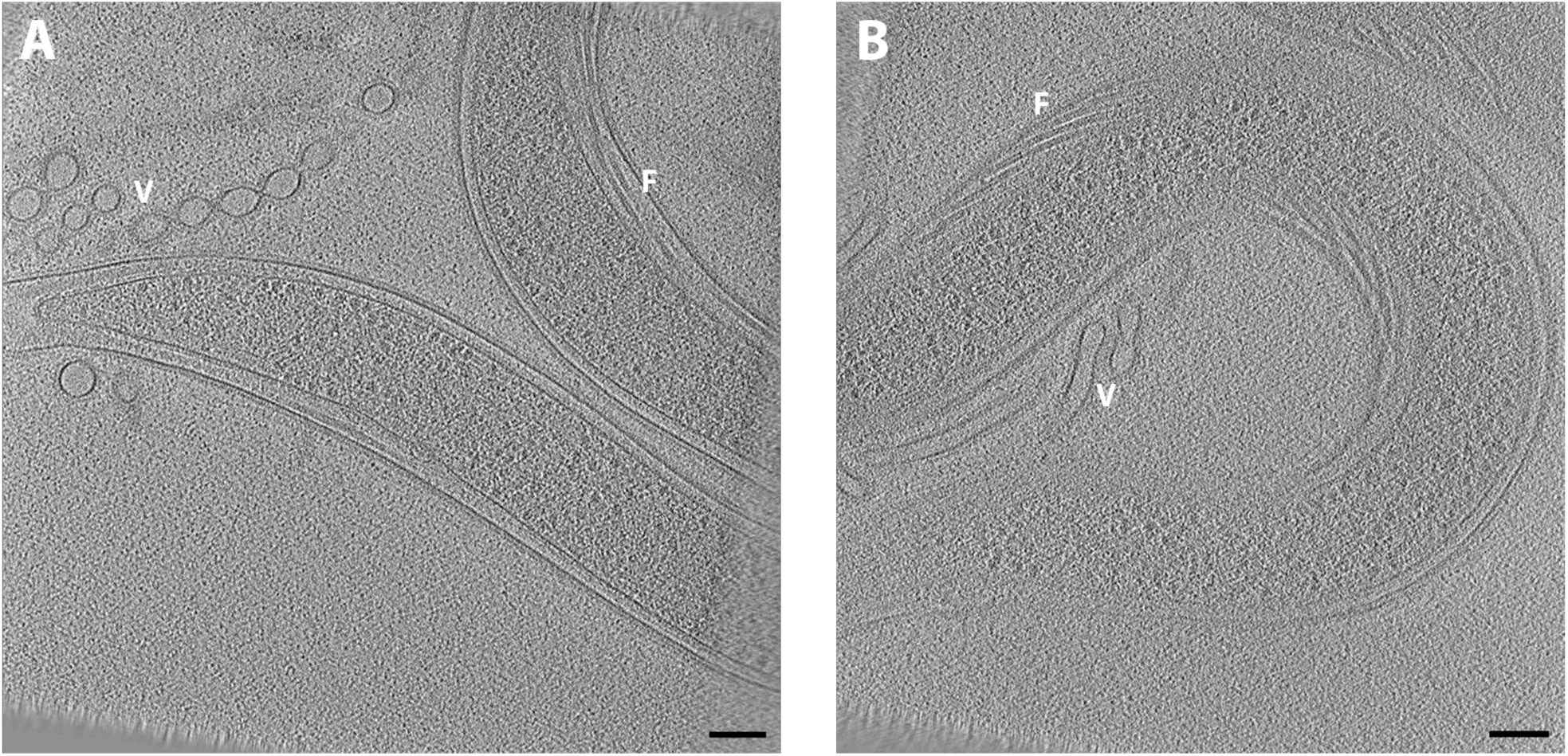
Vesicular outer membrane chains in *Borrelia burgdorferi.* Central tomographic slices of two different *B. burgdorferi* cells illustrate the chained vesicle (**A**) and tubular (**B**) morphologies of membrane-derived structures. V indicates vesicles; F indicates periplasmic flagella. Scale bars 100 nm.

### Intracellular structures

#### “Nanospheres”

In three *Vibrio cholerae* cells (two from a C6706 lacZ^-^ strain (Cameron et al., 2008), and one from a ΔctxA ΔtcpB strain (Chang et al., 2016)) we observed clusters of “nanospheres” – hollow granules with thick walls (Figure 5). The diameter of the nanospheres ranged from ~18-37 nm, and the walls were ~4-10 nm thick. They were pleomorphic: most were roughly spherical, but some were oblong or comma shaped. Each cluster contained about two dozen nanospheres. The clusters were observed at the cell periphery, near the inner membrane (although the clusters were large enough to extend to the center of the cell), and were always observed in close proximity to a filament array structure (discussed below).

**Figure 5.**
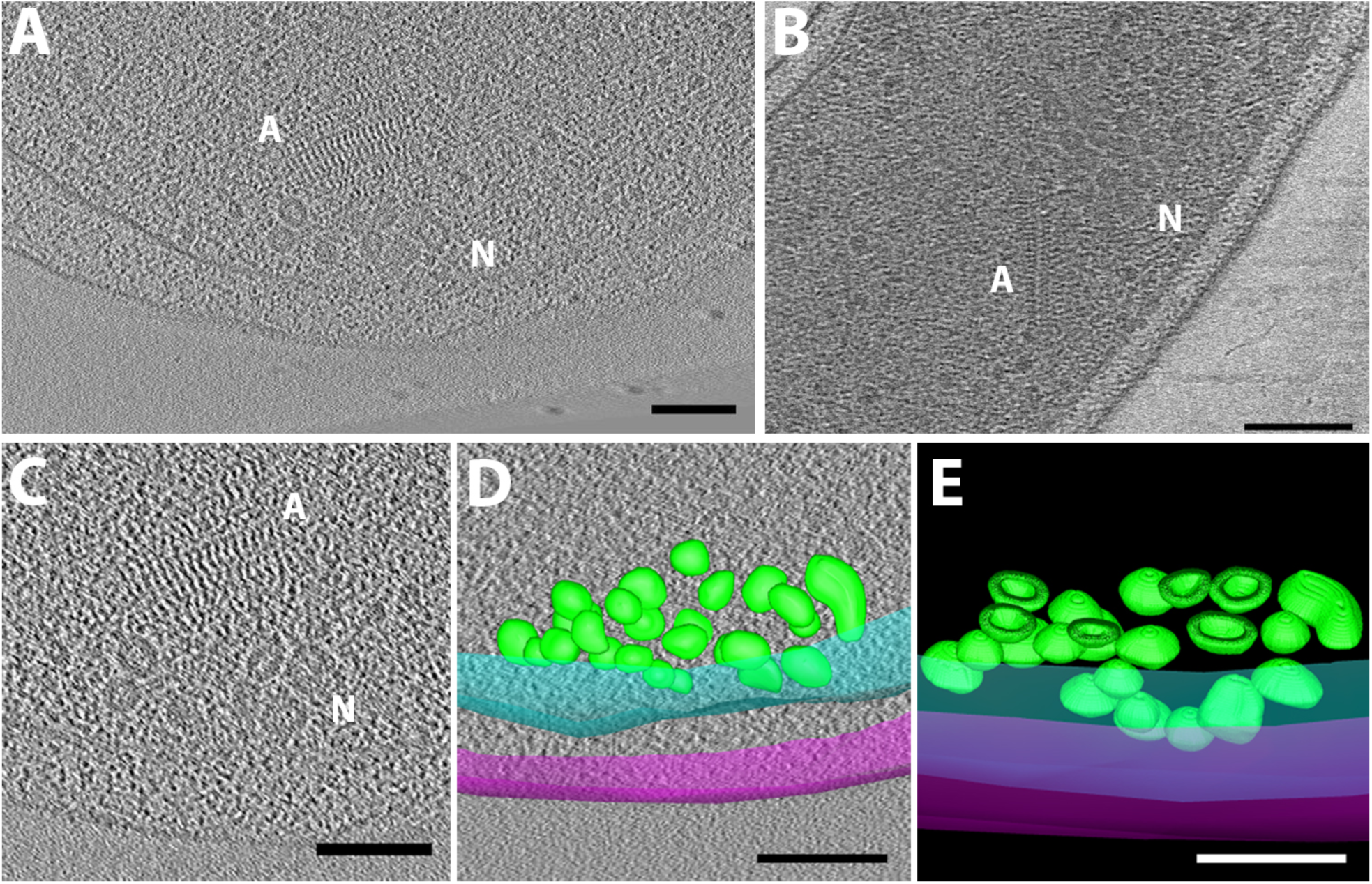
Novel *Vibrio cholerae* nanospheres. Clusters of “nanospheres” were observed in three cryotomograms of *V. cholerae* cells (central slices shown in **A-C**). N indicates nanospheres; A indicates associated filament array. (**D**) shows a segmentation of the cluster seen in (C) overlaid on the tomographic slice, with outer and inner membranes in magenta and cyan, respectively, and nanospheres in green. (**E**) shows a clipping plane through the 3D segmentation revealing the thick walls and hollow centers of the nanospheres. Scale bars 100 nm.

#### Filaments, bundles, arrays, chains and meshes

One of the strengths of ECT imaging is its power to resolve cytoskeletal elements in small bacterial cells. In addition to those we have already identified, we observed many novel filamentous structures in tomograms, including filament arrays, bundles, chains and meshes (Figure 6). In *Hyphomonas neptunium*, we observed long helical filament bundles in the prosthecae that connect dividing cells (Figure 6A). The helix width was 9.5 ± 1.5 nm, the spacing between cross-densities 6.0 ± 0.3 nm, and the helical pitch ~26°. *H. neptunium* divides by asymmetric budding (Weiner et al., 2000) and the genome of the parent cell is passed to the daughter cell through the narrow prostheca connecting the two cells (Zerfas et al., 1997). We observed that the helical structure was straightened in cells treated with ethidium bromide (an intercalator known to unwind DNA (Pommier et al., 1987)) (Figure 6B). We therefore propose that the helix is composed of supercoiled DNA, with each visible filament a DNA duplex connected to adjacent duplexes by cross-densities formed by an unidentified protein.

**Figure 6.**
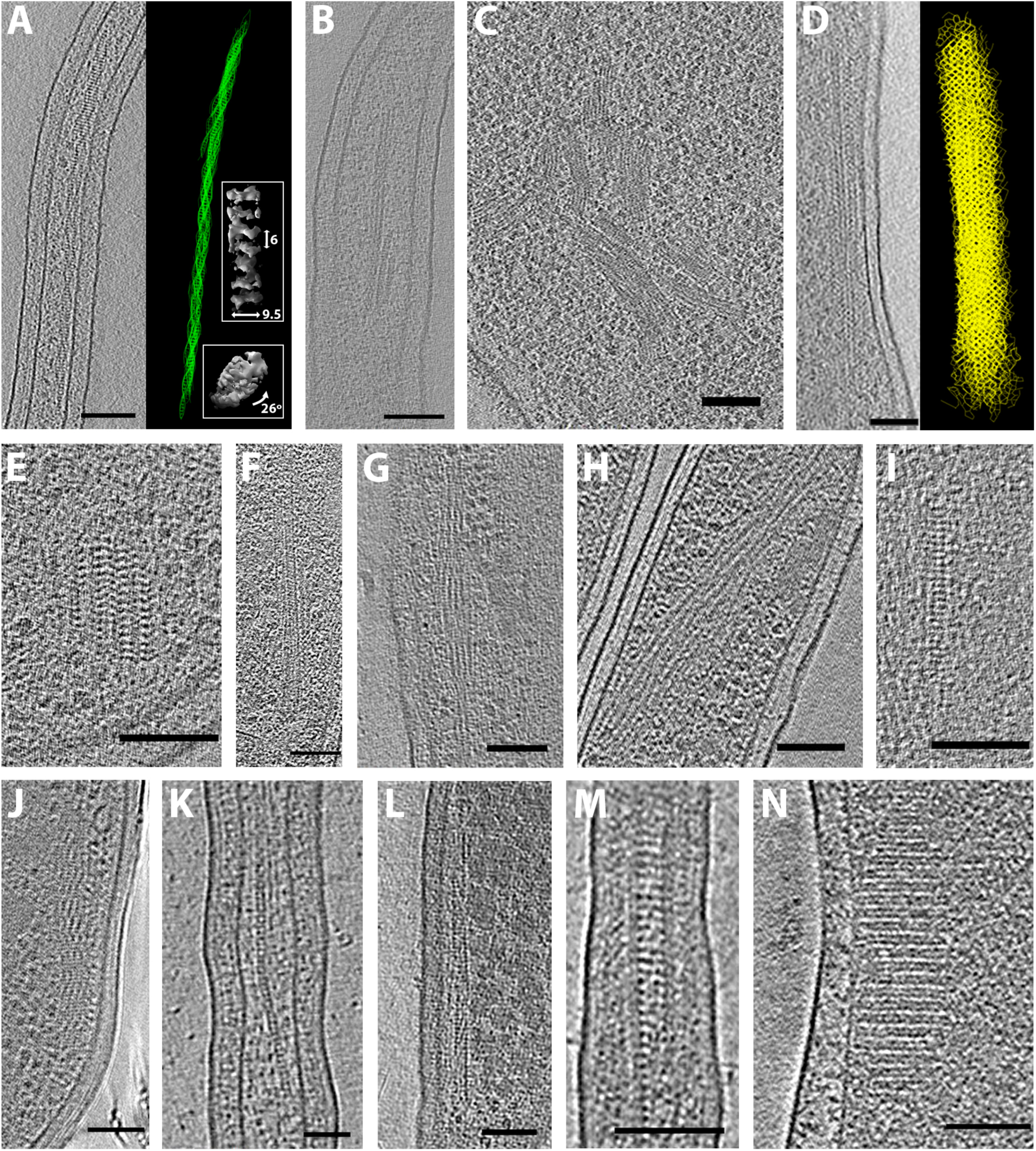
Filament bundles, arrays, and chains. *Hyphomonas neptunium* division stalks contained helical bundles (**A**) that straightened when cells were treated with ethidium bromide (**B**). The right side of panel (**A**) shows a 3D segmentation of the helical bundle, with side and top views of subtomogram averaged insets. Labeled dimensions are in nanometers. (**C**) Large filament bundles in *Helicobacter pylori*. (**D**) A long mesh-like filament array in *Vibrio cholerae*, with segmentation at right. (**E**) A more typical *V. cholerae* filament array. Filament arrays in *Thiomonas intermedia* (**F**), *Hyphomonas neptunium* (**G**) *Hylemonella gracilis* (**H**), *Halothiobacillus neapolitanus* c2 (**I**), and *Mycobacterium smegmatis* (**J**). (**K**) A chain in *Prosthecobacter vanneervenii*. (**L-M**) Filament arrays in *Prosthecobacter debontii*. (**N**) A filament array in a starved *Campylobacter jejuni* cell. Scale bars 100 nm (A-B, D-J, L-N) and 50 nm (C,K).

In *Helicobacter pylori* cells we observed extensive filament bundles. In one cell in an early stage of lysis, such bundles were observed throughout most of the cell (Figure 6C). In *V. cholerae* we observed filament arrays resembling a honeycombed mesh (Figure 6D). These arrays varied in length, but were usually fairly short (~100 nm in length and width), like the example shown in Figure 6E. This is the structure we observed near the nanosphere clusters. Filament arrays exhibited different morphologies in other species. *Thiomonas intermedia* cells contained untwisted arrays ~48 nm thick, ~30 nm wide (Figure 6F). In addition to the prosthecal helix described above, *H. neptunium* cells also contained a bundle of twisting filaments laddered by cross-densities (Figure 6G). These bundles were ~40 nm thick and ~75 nm wide. In a *Hylemonella gracilis* cell we observed a helical bundle of filaments that varied in width and could be related to the nucleoid (Figure 6H). In *Halothiobacillus neapolitanus* c2 cells grown in limited CO_2_ for several hours we observed linear filament arrays with prominent cross-densities spaced 7 ± 0.8 nm apart (Figure 6I). *Mycobacterium smegmatis* displayed straight arrays ~80 nm thick and wide, comprising segments of pitched filaments (Figure 6J). Filament arrays were also seen in multiple species of *Prosthecobacter*: *P. vanneervenii* contained linear chains (Figure 6K) and one *P. debontii* cell contained a straight array similar to those observed in *T. intermedia* (Figure 6L) as well as mesh-like arrays spanning the width of the prostheca (Figure 6M).

In starving *Campylobacter jejuni* cells, we observed regular filament arrays (Figure 6N). When subjected to environmental or cellular stress some bacteria, including *Escherichia coli* have been shown to reorganize their DNA into protective crystalline arrays (Wolf et al., 1999). Since then, additional nucleoid associated proteins have been identified that organize DNA into higher order structures in stationary phase or stress conditions (Teramoto et al., 2010, Lim et al., 2013). The structures we observed in *C. jejuni* resemble those seen in *E. coli* cells overexpressing the protective DNA binding protein Dps (Wolf et al., 1999) and may therefore represent such a nucleoprotein array.

Several other proteins have been shown to copolymerize with DNA into filaments for various functions, including RecA (homologous recombination) (Egelman & Stasiak, 1986) and MuB (DNA transposition) (Mizuno et al., 2013). The width of such filaments *in vitro* (~10 nm) is similar to widths we observed in cells; it is possible that some of the structures in Figure 6 represent these DNA-related processes. Other bacterial proteins form filaments to regulate their function, and it has been suggested that this property may have been coopted in the evolution of the cytoskeleton (Barry & Gitai, 2011). We previously observed such filaments of CTP synthase in tomograms of *Caulobacter crescentus* cells (Ingerson-Mahar et al., 2010). Another protein, alcohol dehydrogenase, forms plaited filaments ~10 nm wide, called spirosomes, in many bacteria capable of anaerobic metabolism (Matayoshi et al., 1989, Laurenceau et al., 2015). It is possible that some of the filament arrays and chains we observed in tomograms may be filaments formed by these or other, yet uncharacterized, proteins.

In addition to filament arrays and bundles, we observed individual or paired filaments in nearly every species imaged. Examples are shown in Figure 7. (Note that due to their ubiquity, statistics are not included in Supplementary Table 1.) Filaments were seen with various orientations in the cytoplasm (Figure 7A-C), as well as running alongside the membrane (Figure D-E). Consistent with our previous work (Swulius et al., 2011), we did not observe any arcing filaments immediately adjacent to the membrane as predicted by some studies of MreB (e.g. (Jones et al., 2001, Shih et al., 2003)). (Note that we did observe filaments corresponding to the known types of MamK (Komeili et al., 2006, Scheffel et al., 2006), FtsZ (Li et al., 2007, Szwedziak et al., 2014), and bactofilins (Kuhn et al., 2010) but we do not show them here since they have already been characterized.) Paired filaments have been shown to function in plasmid segregation, so it is possible that some paired filaments we observed were such ParM or TubZ structures (Aylett et al., 2010, Bharat et al., 2015).

**Figure 7.**
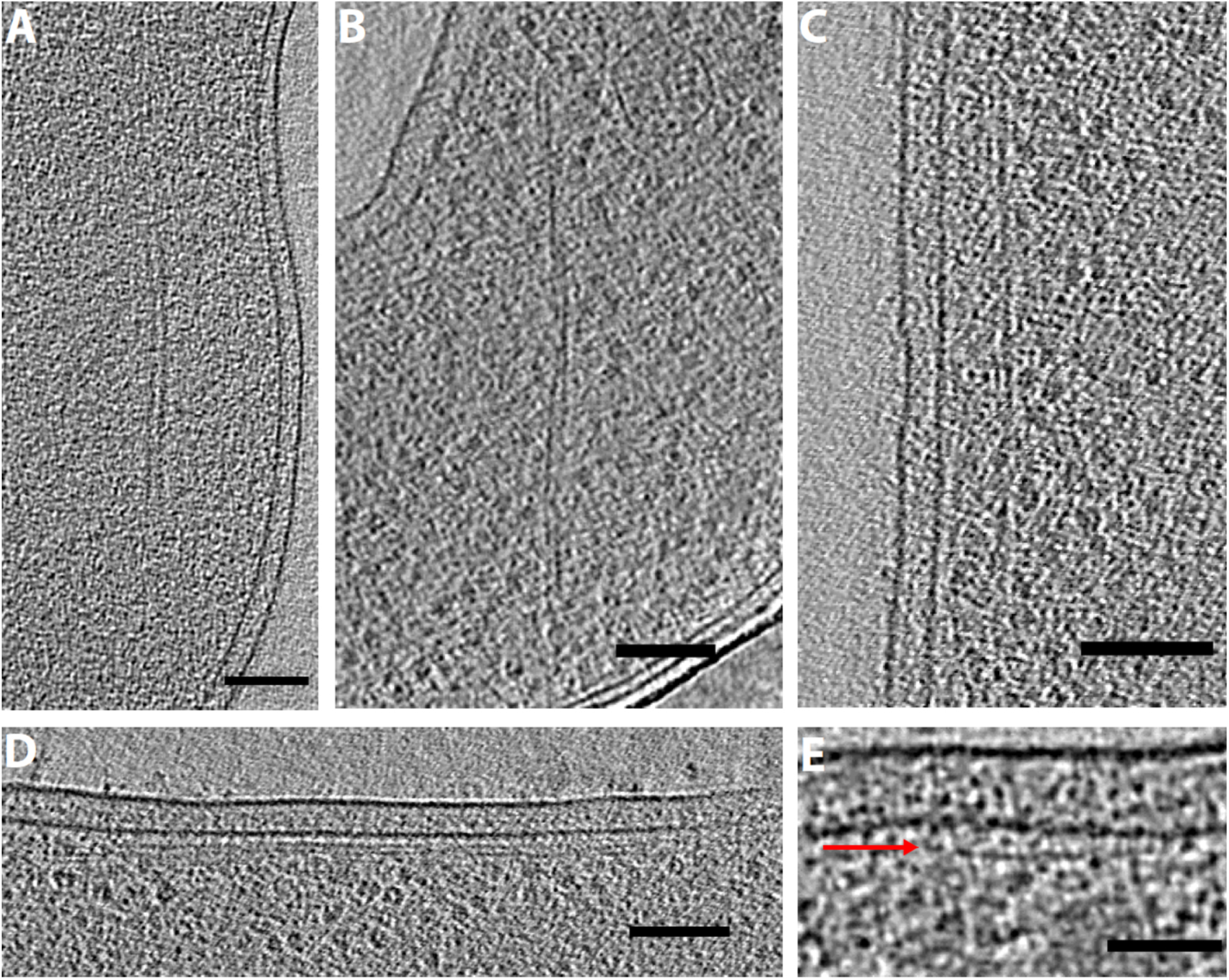
Single and paired filaments. Tomographic slices showing paired filaments in *Campylobacter jejuni* (**A**), and *Thiomicrospira crunogena* (**B**), and membrane-aligned filaments in *Shewanella putrefaciens* (**C**), *Prosthecobacter debontii* ***(D)***, and *Prosthecobacter fluviatilis* (**E**, red arrow shows filament just under the inner membrane). Scale bars 100 nm (A-D) and 50 nm (E).

#### Tubes

In addition to the known types of tubes we have reported earlier (bacterial microtubules (Pilhofer et al., 2011) and type VI secretion systems (Basler et al., 2012)), we observed several novel tubular structures in bacterial cells (Figure 8). In *Thiomicrospira crunogena* we found large tubes (18.6 ± 1.8 nm diameter) containing eight outer protofilaments surrounding a central protofilament (Figure 8A). *H. neapolitanus* c2 cells also contained large tubes (16.7 ± 0.7 nm diameter) with a central filament (Figure 8B). In several other species, we observed hollow tubes of varying dimensions: 8.9 ± 0.3 nm diameter in *Bdellovibrio bacteriovorus* (Figure 8C), 14.3 ± 1.7 nm in *T. intermedia* (Figure 8D), and 8.3 ± 0.5 nm in *H. neptunium* (Figure 8E). *H. neptunium* cells also contained many rings of similar diameter. In fact, we observed rings in many species, which could be assembly or disassembly intermediates of tubes.

**Figure 8.**
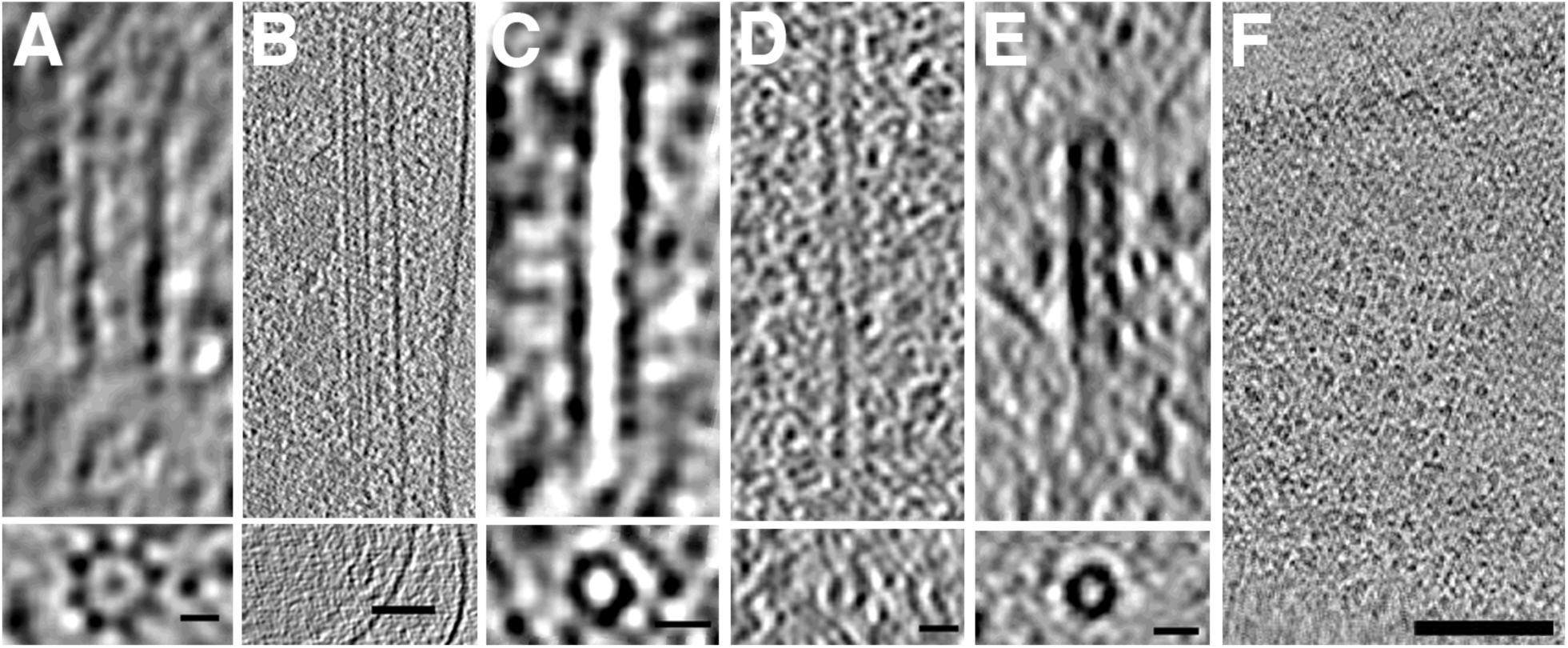
Tubes and rings. Tubes observed in *Thiomicrospira crunogena* (**A**), *Halothiobacillus neapolitanus c2* (**B**), *Bdellovibrio bacteriovorus* (**C**), *Thiomonas intermedia* (**D**), and *Hyphomonas neptunium* (**E**). In each panel, tomographic slices show a side view (above), and a top view (bottom). (**F**) An array of rings observed in *Helicobacter pylori*. Scale bars 10 nm (A,C,E), 20 nm (B,D), and 100 nm (F).

In addition to isolated rings, in one case we observed an organized array of rings. One slightly lysed (a condition that flattens the cell and increases image quality) *H. pylori* cell contained a striking array of about two dozen evenly spaced rings near the cytoplasmic membrane (Figure 8F). Each ring was ~6 nm in diameter and ~20 nm (center-to-center distance) from its neighbors in the square lattice.

#### Vesicles

In contrast to eukaryotic cells, relatively little is known about membrane remodeling in Bacteria. Compartmentalized cells in the Planktomycetes-Verrucomicrobia-Chlamydiae (PVC) superphylum have been shown to contain homologs of eukaryotic membrane trafficking proteins (Santarella-Mellwig et al., 2010) and exhibit endocytosis-like protein uptake (Lonhienne et al., 2010). An additional potential membrane-remodeling system based on FtsZ homologues is more widespread across Bacteria, but its function remains unknown (Makarova & Koonin, 2010).

Despite this limited evidence for membrane remodeling in Bacteria, we observed intracellular vesicles in nearly every species imaged. They exhibited various sizes, shapes, membrane layers, and contents. They were frequently found near the cytoplasmic membrane. Figure 9 shows examples of round and horseshoe-shaped vesicles. Round vesicles were found in nearly every species imaged, and therefore no statistics for them are compiled in Supplementary Table 1. Most round vesicles were empty (density similar to background; e.g. Figure 9A-C). One of these vesicles, observed in a lysed cell (improving clarity by reducing cytoplasmic crowding), exhibited regularly spaced protein densities around its exterior (Figure 9C). Others were at least partially filled with denser material (e.g. Figure 9D-F). In two *M. xanthus* cells overexpressing a fluorescent fusion of a periplasmic protein (PilP-sfGFP), we observed round vesicles containing a dense amorphous core (Figure 9F). These could be a novel form of membrane-bound inclusion body, perhaps packaged from the periplasm. In eight species, we observed horseshoe-shaped vesicles (Figure 9G-H).

**Figure 9.**
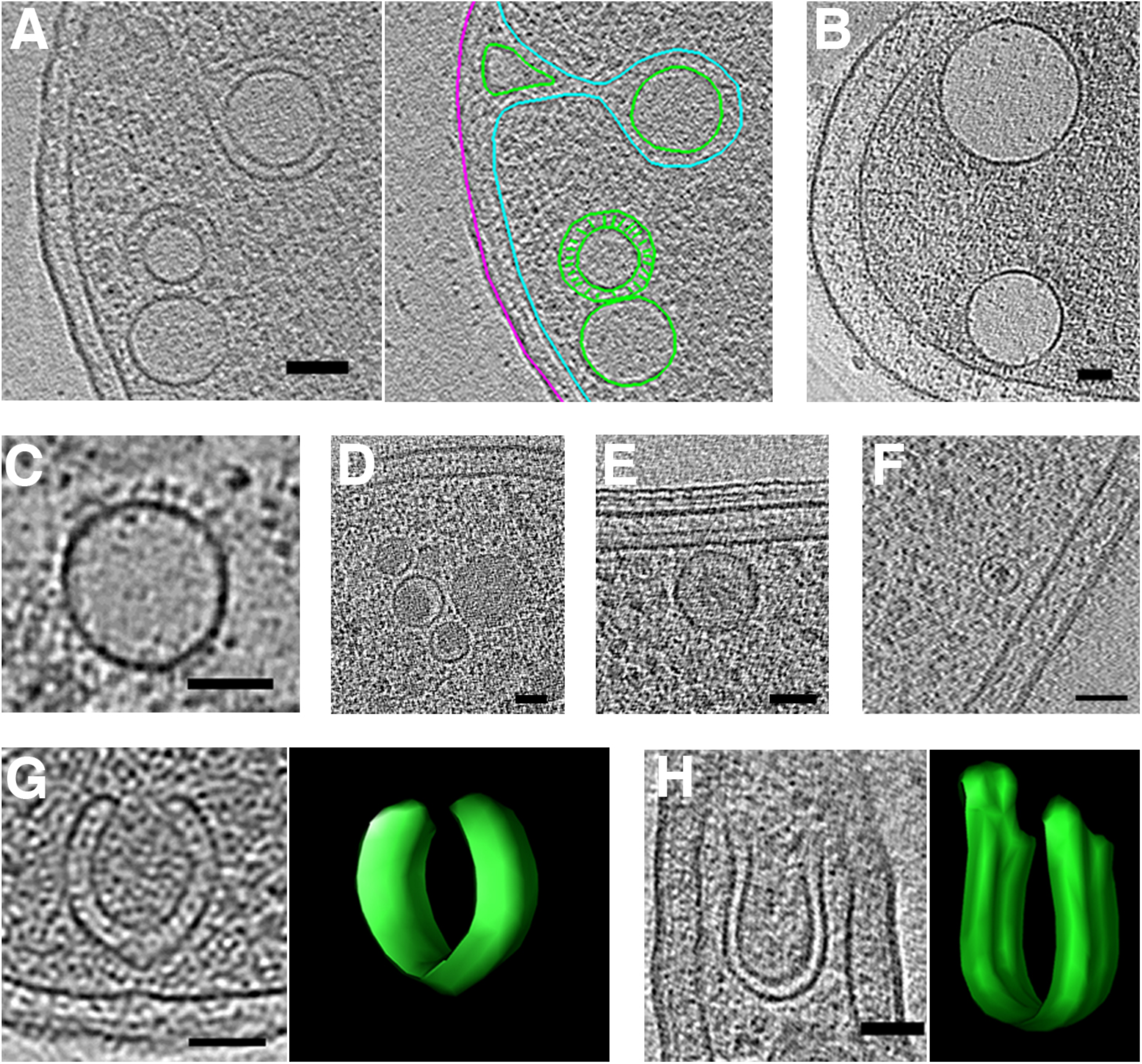
Round and horseshoe-shaped vesicles. Tomographic slices showing examples of round vesicles in *Escherichia coli* (**A**, segmentation shown at right), *Helicobacter pylori* (**B**), *Helicobacter hepaticus* (**C**), *Myxococcus xanthus* (**D**), *Caulobacter crescentus* (**E**), and *Myxococcus xanthus* overexpressing PilP-sfGFP (**F**). Examples of horseshoe-shaped vesicles in *Ralstonia eutropha* (**G**) and *Prosthecobacter fluviatilis* (**H**), with 3D segmentations shown at right. In the segmentation in (A), outer and inner membranes are in magenta and cyan, respectively, and vesicles in green. Scale bars 50 nm.

Flattened vesicles (Figure 10A-F) were less common than round vesicles, and were usually observed near membranes or wrapping around storage granules (Figure 10E), suggesting a possible functional relationship. Flattened vesicles were usually empty. One *T. intermedia* cell contained a stack of flattened vesicles (Figure 10A). Flattened vesicles were particularly prevalent in *C. crescentus* cells (Figure 10B-E). *P. debontii* cells contained flattened vesicles that neither ran along the membrane nor wrapped around granules (Figure 10F). Since the lowest energy shape of a liposome is a sphere, it is likely that the vesicles were flattened by cytoplasmic pressure or some other constraint such as associated protein.

**Figure 10.**
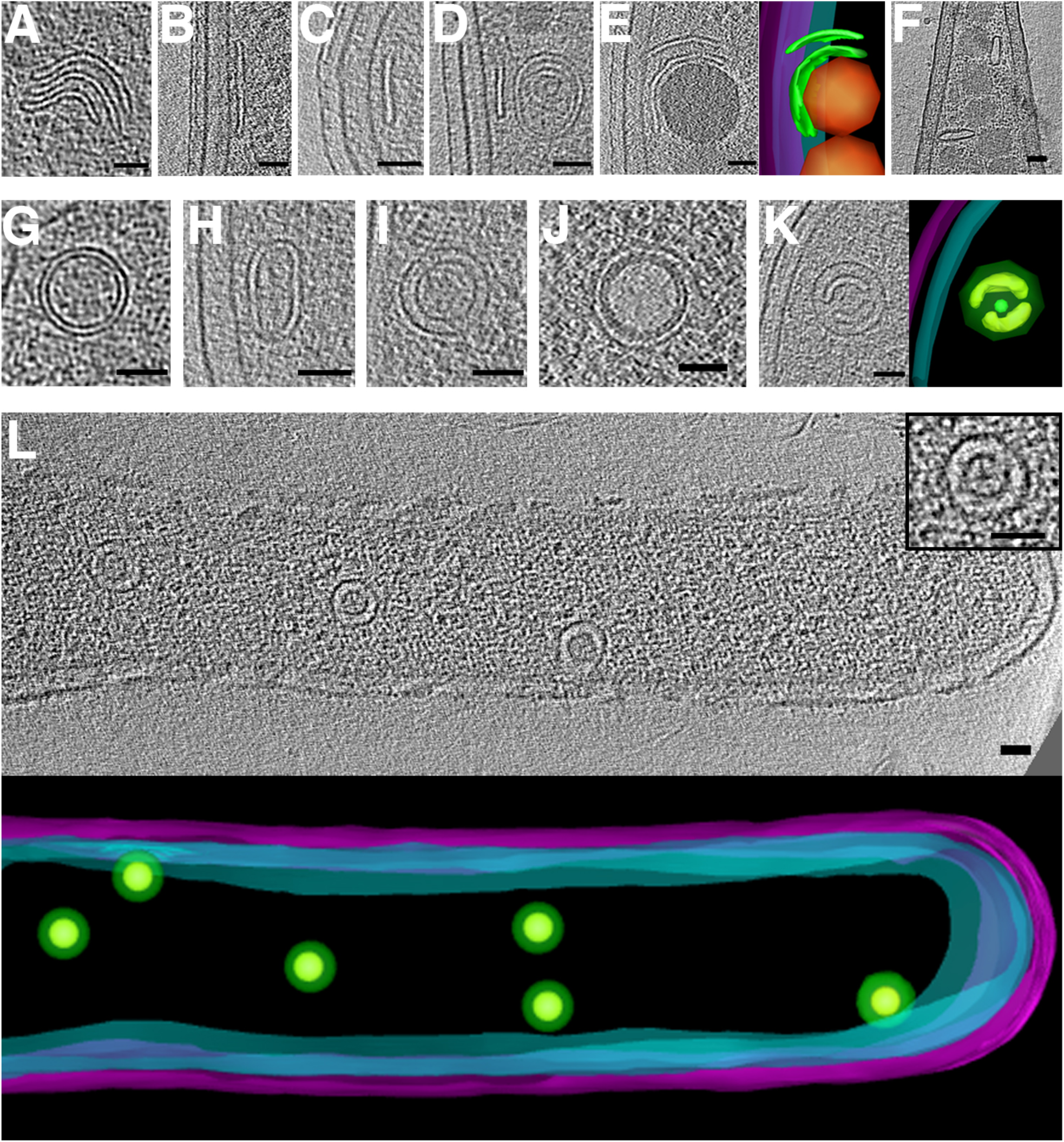
Flattened and nested vesicles. Examples of flattened vesicles in *Thiomonas intermedia* (**A**), *Caulobacter crescentus* (**B-E**) and *Prosthecobacter debontii* (**F**). Note storage granules in (E and F), shown in orange in the segmentation in (E). Examples of nested vesicles in *Serpens flexibilis* (**G**), *Caulobacter crescentus* (**H**), *Borrelia burgdorferi* (**I**), *Vibrio cholerae* (**J**), *Caulobacter crescentus* with segmentation (**K**), and strain JT5 (**L**). Inset in (L) shows an enlargement of central vesicle, and a 3D segmentation of the visible portion of the cell is shown below. In segmentations, outer and inner membranes are shown in magenta and cyan, respectively, and vesicles in green. Scale bars 50 nm.

Many cells contained nested vesicles, with diverse sizes and shapes, as well as subcellular locations (Figure 10G-L). In some nested vesicles, densities were observed bridging the inner and outer membranes (Figure 10G-H). Cells of strain JT5 exhibited multiple nested vesicles of uniform shape and size (Figure 10L).

We also observed periplasmic vesicles in many species (Figure 11). They were typically empty and exhibited great variability in size, shape, and abundance. In some cases, they were even seen to form branching networks (Figure 11A). As with cytoplasmic vesicles, they were most abundant in cells showing signs of stress.

**Figure 11.**
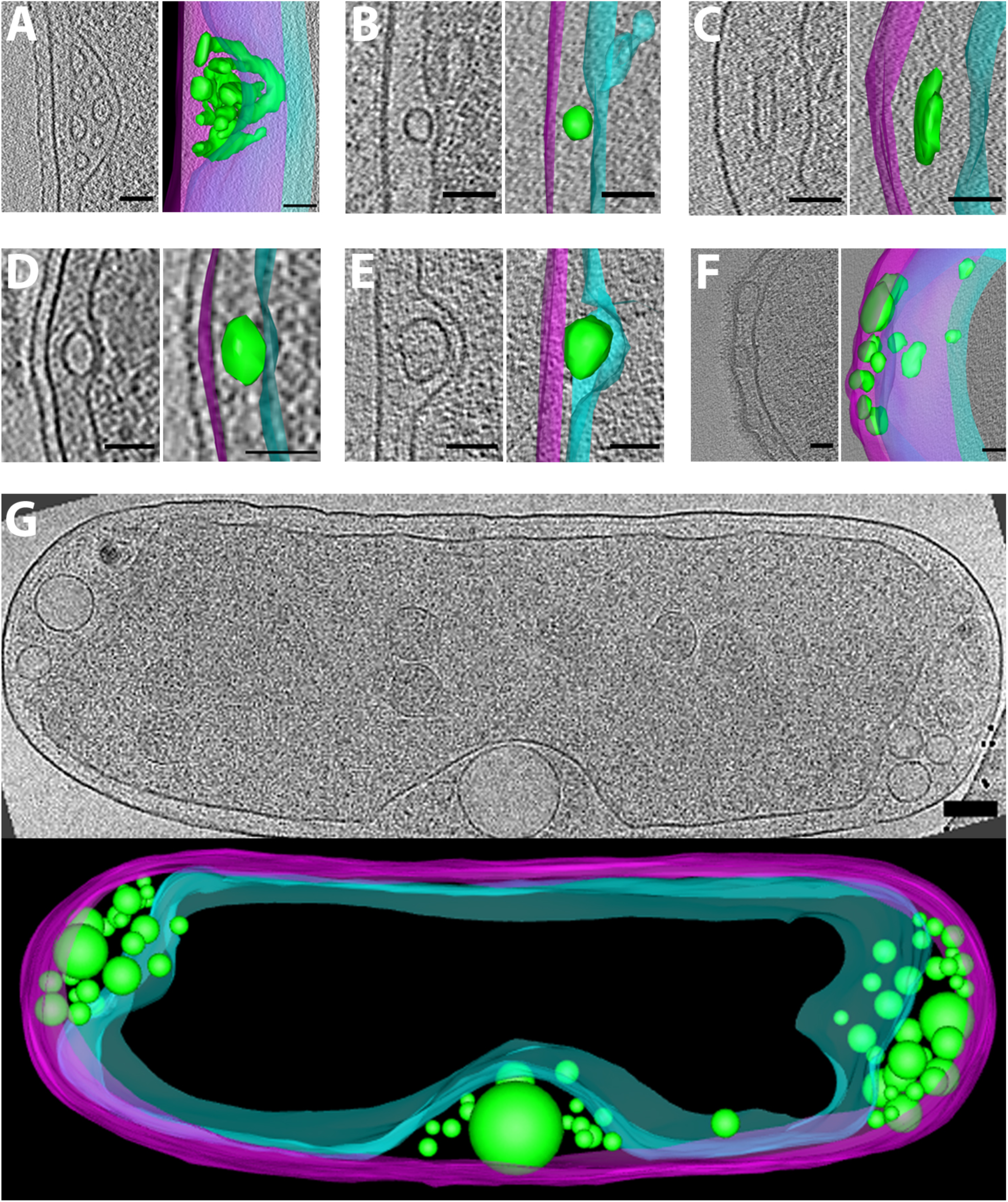
Periplasmic vesicles. Examples of periplasmic vesicles in *Caulobacter crescentus* (**A**), *Helicobacter pylori* (**Β**), *Brucella abortus* (**C**), *Thiomonas intermedia* (**D**), *Hyphomonas neptunium* (**E**), *Myxococcus xanthus* (**F**), and *Halothiobacillus neapolitanus* c2 (**G**). In each panel, a central tomographic slice is shown, as well as a segmentation with outer and inner membranes in magenta and cyan, respectively, and vesicles in green. Scale bars 50 nm (A-F) and 100 nm (G).

## Conclusions

Here we present the results of a survey of, to our knowledge, uncharacterized bacterial structures that we have observed in our work over the last 10+ years. We hope that further study will identify them and their functions. Already, they signal the wealth of complexity still to be discovered in bacterial cells.

## MATERIALS and METHODS

### Strains and growth

Unless otherwise noted, bacterial strains were wild-type and grown in species-standard medium and conditions to mid-log or early stationary phase. *Azospirillum brasilense* cultures were switched to nitrogen-free medium for ~16 hours prior to imaging to induce nitrogen fixation and digestion of storage granules that decrease image quality. Predatory *Bdellovibrio bacteriovorus* cells were co-cultured with *Vibrio cholerae* strain MKW1383. *Helicobacter pylori* cells were cultured with human gastric carcinoma cells. In all cases, samples of cells in growth medium were mixed with BSA-treated 10nm colloidal gold fiducials (Sigma), applied to glow-discharged EM grids (Quantifoil), and plunge-frozen in a liquid ethane-propane mixture (Tivol et al., 2008). Grids were maintained at liquid nitrogen temperature throughout storage, transfer, and imaging.

### Electron cryotomography

Plunge-frozen samples were imaged using either a Polara or Titan Krios 300 kV FEG transmission electron microscope (FEI Company) equipped with an energy filter (Gatan). Images were recorded using either a lens-coupled 4k x 4k UltraCam CCD (Gatan) or a K2 Summit direct electron detector (Gatan). Tilt-series were recorded from −60° to +60° in 1-2°increments, with defoci of ~6-12 μm and a cumulative dose of ~100-200 e^-^/Å^2^. Tilt-series were acquired automatically using either Leginon (Suloway et al., 2009) or UCSF Tomography (Zheng et al., 2007) software. Tomographic reconstructions were calculated using either the IMOD software package (Kremer et al., 1996) or Raptor (Amat et al., 2008). 3D segmentations and movies were produced with IMOD (Kremer et al., 1996). Subtomogram averages were calculated using PEET software (Nicastro et al., 2006).

## ACKNOWLEDGMENTS

The authors would like to thank our collaborators who provided strains for imaging: Andrew Camilli (*Streptococcus pneumoniae*), Eric Matson (strain JT5), Gladys Alexandre (*Azospirillum brasilense* mutants), Lotte Søgaard-Andersen, Simon Ringgaard and Matthew K. Waldor (*Vibrio cholerae* wild type and mutants), Michael Marletta (*Shewanella putrefaciens*), and Gordon Cannon and Sabine Heinhorst (*Halothiobacillus neapolitanus and Thiomonas intermedia*). We also thank members of the Jensen lab for helpful discussions.

### COMPETING INTERESTS

No competing interests declared.

### AUTHOR CONTRIBUTIONS

Conceptualization: M.J.D. and G.J.J.; formal analysis and investigation: M.J.D., C.M.O., A.P., J.C., K.G., T.J., J.T., J.D., Y.P., A.K., A.I.J., M.P., S.C., E.I.T., Y.-W.C., A.B., J.S., Z.L., P.S., C.V.I., B.A.S., A.W.M.; writing – original draft preparation: M.J.D. and C.M.O.; writing – review and editing: M.J.D., C.M.O., G.J.J.; funding acquisition: M.J.D. and G.J.J.; resources: G.J.J.; supervision: M.J.D. and G.J.J.

## Supplementary Movies 1-11

The movies show all figure panels going slice-by-slice through the Z-stack of the tomogram. Accompanying models are provided when a three-dimensional view is helpful for interpreting the structure. https://figshare.com/s/782461843c3150d27cfa

**Supplementary Table 1.** Species range and frequency of structures observed. For species with many tomograms imaged we are reporting the lowest estimate for counts since not all tomograms were viewed in search of each feature.

| Feature | Species | # Tomograms | % Tomograms |
| --- | --- | --- | --- |
| Surface appendages | *Prostheco bacter debontii* | 6 | 4.6 |
| Surface hooks | *Azospirillum brasilense* WT | 20 | 95.2 |
| | *A. brasilense* ΔcheOp1 | 24 | 92.3 |
| | *A. brasilense* ΔcheOp4 | 9 | 52.9 |
| | *A. brasilense* ΔcheOp1 ΔcheOp4 | 0 | 0.0 |
| Polar fimbriae | Strain JT5 (*Dysgonomonas*) | 41 | 100 |
| OMV chains | *Borrelia burgdorferi* | 24 | 52.2 |
| Nanospheres | *Vibrio cholerae* | 3 | 0.2 |
| Filament bundles | *Agrobacterium tumefaciens* | 1 | < 0.1 |
|  | *Borrelia burgdorferi* | 1 | 2.2 |
|  | *Campylobacter jejuni* | 2 | 3.3 |
|  | *Caulobacter crescentus* | 1 | < 0.1 |
|  | *Escherichia coli* | 1 | 0.1 |
|  | *Halothiobacillus neapolitanus* c2 | 5 | 2.6 |
|  | *Helicobacter pylori* | 2 | 2.1 |
|  | *Hylemonella gracilis* | 1 | 0.9 |
|  | *Hyphomonas neptunium* | 32 | 2.7 |
|  | *Mycobacterium smegmatis* | 5 | 5.8 |
|  | *Myxococcus xanthus* | 1 | 0.1 |
|  | *Prostheco bacter debontii* | 2 | 1.5 |
|  | *Prostheco bacter fluviatilis* | 5 | 3.6 |
|  | *Prostheco bacter vanneervanii* | 4 | 2.3 |
|  | *Thiomonas intermedia* | 3 | 6.7 |
|  | *Vibrio cholerae* | 10 | 0.8 |
|  | *Shewanella putrefaciens* | 1 | 6.3 |
| Tubes | *Bdellovibrio bacteriovorus* | 1 | 0.3 |
|  | *Halothiobacillus neapolitanus* c2 | 1 | 0.5 |
|  | *Hyphomonas neptunium* | 7 | 0.6 |
|  | *Prostheco bacter vanneervanii* | 1 | 0.6 |
|  | *Thiomicrospira crunogena* | 2 | 9.1 |
|  | *Thiomonas intermedia* | 1 | 2.2 |
| Ring array | *Helicobacter pylori* | 1 | 1.0 |
| Vesicles - horseshoe | *Caulobacter crescentus* | 8 | 0.4 |
|  | *Hyphomonas neptunium* | 1 | 0.1 |
|  | *Magnetospirillum magneticum* | 1 | 0.2 |
|  | *Myxococcus xanthus* | 2 | 0.1 |
|  | *Prostheco bacter fluviatilis* | 2 | 1.4 |
|  | *Prostheco bacter vanneervanii* | 1 | 0.6 |
|  | *Ralstonia eutropha* | 1 | 1.6 |
|  | *Thiomonas intermedia* | 2 | 4.4 |
| Vesicles - flattened | *Agrobacterium tumefaciens* | 1 | < 0.1 |
|  | *Caulobacter crescentus* | 78 | 3.6 |
|  | *Escherichia coli* | 1 | 0.1 |
|  | *Helicobacter pylori* | 16 | 2.9 |
|  | *Hyphomonas neptunium* | 1 | 0.1 |
|  | *Myxococcus xanthus* | 9 | 0.5 |
|  | *Prostheco bacter debontii* | 1 | 0.8 |
|  | *Prostheco bacter dejongeii* | 7 | 5.3 |
|  | *Prostheco bacter fluviatilis* | 4 | 2.9 |
|  | *Prostheco bacter vanneervanii* | 1 | 0.6 |
|  | *Streptococcus pneumoniae* | 4 | 11.4 |
|  | *Thiomonas intermedia* | 1 | 2.2 |
|  | *Treponema primitia* | 1 | 1.1 |
|  | *Vibrio cholerae* | 3 | 0.2 |
| Vesicles - nested | *Agrobacterium tumefaciens* | 2 | 0.1 |
|  | *Bacillus subtilis* | 1 | 1.9 |
|  | *Bdellovibrio bacteriovorus* | 1 | 0.3 |
|  | *Borrelia burgdorferi* | 1 | 2.2 |
|  | *Brucella abortus* | 2 | 3.7 |
|  | *Caulobacter crescentus* | 53 | 2.4 |
|  | *Escherichia coli* | 4 | 0.3 |
|  | *Halothiobacillus neapolitanus* c2 | 5 | 2.6 |
|  | *Helicobacter hepaticus* | 2 | 6.1 |
|  | *Helicobacter pylori* | 10 | 1.8 |
|  | *Hyphomonas neptunium* | 3 | 0.3 |
|  | Strain JT5 (*Dysgonomonas*) | 2 | 4.7 |
|  | *Myxococcus xanthus* | 23 | 1.3 |
|  | *Prostheco bacter debontii* | 3 | 2.3 |
|  | *Prostheco bacter dejongei* | 3 | 2.3 |
|  | *Prostheco bacter fluviatilis* | 1 | 0.7 |
|  | *Prostheco bacter vanneerveni* | 2 | 1.1 |
|  | *Ralstonia eutropha* | 2 | 3.2 |
|  | *Serpens flexibilis* | 1 | 10.0 |
|  | *Sphingopyxis alaskensis* | 1 | 4.5 |
|  | *Streptococcus pneumoniae* | 1 | 2.9 |
|  | *Tetrasphaera remis* | 1 | 25.0 |
|  | *Thiomonas intermedia* | 1 | 2.2 |
| Vesicles - periplasmic | *Agrobacterium tumefaciens* | 2 | 0.1 |
|  | *Bdellovibrio bacteriovorus* | 2 | 0.6 |
|  | *Brucella abortus* | 3 | 5.6 |
|  | *Campylobacter jejuni* | 1 | 1.7 |
|  | *Caulobacter crescentus* | 16 | 0.7 |
|  | *Escherichia coli* | 6 | 0.5 |
|  | *Halothiobacillus neapolitanus* | 22 | 11.5 |
|  | *Helicobacter hepaticus* | 2 | 6.1 |
|  | *Helicobacter pylori* | 3 | 0.5 |
|  | *Hyphomonas neptunium* | 3 | 0.3 |
|  | *Myxococcus xanthus* | 1 | 0.1 |
|  | *Prostheco bacter dejongei* | 1 | 0.8 |
|  | *Prostheco bacter fluviatilis* | 2 | 1.4 |
|  | *Prostheco bacter vanneerveni* | 3 | 1.7 |
|  | *Ruminococcus flavefaciens* | 2 | 18.2 |
|  | *Thiomicrospira crunogena* | 1 | 4.5 |
|  | *Thiomonas intermedia* | 1 | 2.2 |
|  | *Vibrio cholerae* | 2 | 0.2 |

